# Prefrontal cortex and striatum dissociate modality and learning order in cross-modal sensorimotor learning

**DOI:** 10.1101/2025.11.10.687671

**Authors:** Da Song, Andrew J. Peters

## Abstract

Sensorimotor learning is accompanied by the emergence and strengthening of sensory responses in the prefrontal cortex (PFC) and striatum. However, it remains unclear whether learning-evoked sensory responses in these structures are modality-specific, or whether they represent a more generalized signal for modality-independent but behaviorally relevant stimuli. To address this question, we performed widefield calcium imaging while mice were trained on sensorimotor tasks involving either visual or auditory stimuli, with mice first learning one modality and then switching to the other. We found that the anterior PFC (aPFC) exhibited a bilateral response to both visual and auditory stimuli after learning the respective tasks, indicating a generalized sensory representation. In contrast, the medial PFC (mPFC) showed modality-specific activity, responding only to contralateral visual stimuli after learning. Mice were able to learn sensorimotor associations for both modalities in sequence, and were only able to transfer learned behavior from visual to auditory stimuli but not the reverse. Despite differences in cross-modal learning transfer, cortical responses to learned stimuli were the same regardless of learning order. In contrast, electrophysiological recordings in the anterior striatum revealed that visually responsive regions became co-responsive to auditory stimuli only in mice trained on the visual task first. The additional auditory responses arose from distinct neurons rather than those previously responsive to visual stimuli, indicating that co-responsiveness occurred at the regional rather than single-neuron level. Together, these findings reveal that learned sensory responses in the PFC and striatum uniquely dissociate modality and learning order, suggesting a divergence in their roles during sensorimotor learning.

## Introduction

Learned associations between stimuli and actions are not necessarily one-to-one. Instead, a single behavior can be triggered by one of many different sensory cues. For instance, when crossing a street, walking can be cued by either a visual walk sign or an auditory crossing noise. In this case, sensory streams from distinct modalities converge on a common motor output. However, it is not known whether there are separate sensorimotor pathways for each modality that must be established independently, or if sensory responses converge into multimodal regions which support more generalized sensorimotor associations.

Two key regions for investigating this question are the prefrontal cortex (PFC) and anterior striatum. These regions exhibit increased^2^or novel^3^responses to sensory stimuli after learning a sensorimotor association. This suggests that learning causes sensory information to become routed into these regions, which may act as mediators between sensory and motor areas. It is unclear whether these learned sensory responses are multimodal and generally mark the presence of any behaviorally relevant cue^4-6^, or if there are separate subregions within PFC and anterior striatum that respond to specific modalities^7^.

Previous work has provided evidence for both modality-convergent and modality-parallel pathways in sensorimotor learning. Towards convergent pathways, both the PFC^3,4,8-12^and anterior striatum^2,13,14^are recruited during sensorimotor learning across modalities. Furthermore, inputs from different sensory cortices converge in both the PFC^15,16^and anterior striatum^17-19^, providing an anatomical substrate for multimodal function. Towards parallel pathways, stimulus responses are heterogeneous across subregions of the PFC^15,20,21^, and the functional and anatomical modality overlap in the anterior striatum remains uncertain, as does the role of this overlapping domain compared to the more segregated posterior regions^22-24^.

One approach to distinguish between convergent and parallel architectures is to ask whether a sensorimotor rule learned in one modality can be re-used for another modality. Training animals first on a single modality and then switching to a different modality provides such a test: successful cross-modal transfer would imply that overlapping circuits are engaged, whereas failure of transfer would indicate that a new, modality-specific pathway must be established^25,26^.

We tested this by training mice to associate either a visual or auditory stimulus with a forelimb movement, and then switching mice to the alternate modality. Widefield calcium imaging revealed that the medial PFC (mPFC) developed responses only to task-relevant visual stimuli, whereas the anterior PFC (aPFC) developed responses to relevant stimuli across both modalities. Behaviorally, visuomotor learning transferred to the audiomotor task, but not vice versa, revealing an asymmetric transfer between modalities. The order of learned modalities did not affect post-learning stimulus responses in the cortex, but did result in different stimulus responses in the anterior striatum, suggesting a possible site of cross-modal transfer. Together, these findings uncover a modality- and order-dependence in cortical and striatal sensory responses during sensorimotor learning, with the PFC distinguishing modality but not learning order, and the striatum distinguishing both modality and learning order.

## Results

### Modality-specific regions in the prefrontal cortex

We trained mice to develop a sensorimotor association with either visual or auditory stimuli. Mice were head-fixed in the center of three screens with a centrally located speaker, and rested their forepaws on a wheel that could be turned clockwise and counterclockwise. In both tasks, a stimulus was presented with a property that was yoked to the wheel movement, and counterclockwise wheel movements would trigger a sucrose reward (**Extended Data Fig. 1a**). In the visual variant of the task, the stimulus was a vertical grating displayed on the right-hand screen, and turning the wheel counterclockwise moved the stimulus leftward to trigger a reward once it reached the center (**Fig. 1a**)^3^. In the auditory variant of the task, the stimulus was an 8 kHz pure tone, and turning the wheel counterclockwise lowered the tone volume to trigger a reward once the volume reached zero (**Fig. 1b**).

**Figure 1.**
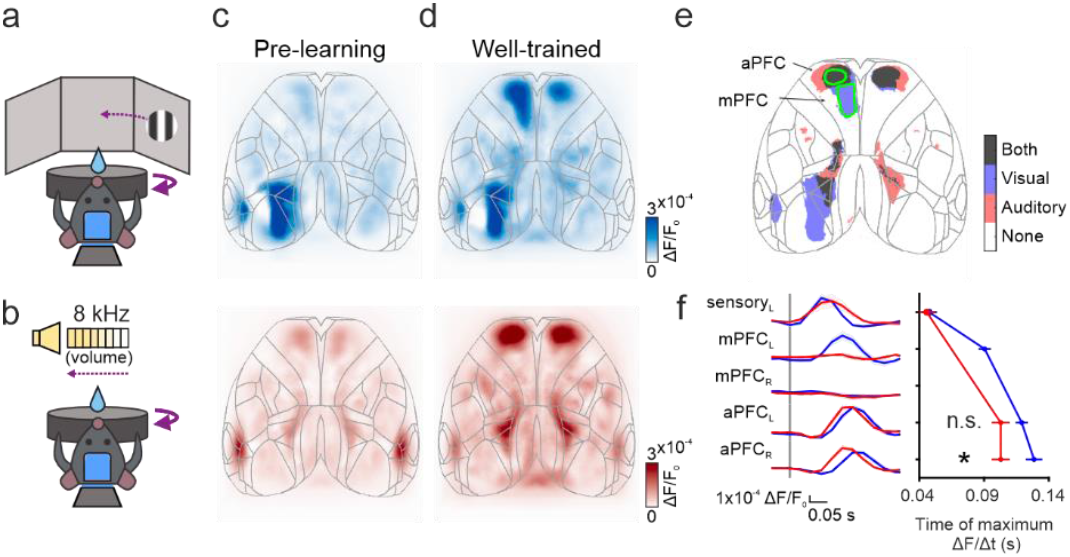
Visuomotor and audiomotor learning induce sensory responses in different subregions of the PFC. **aa-b**, Behavioral paradigm of visuomotor (a) and audiomotor (b) learning task. **c**, Cortical responses to task stimuli before learning, expressed as kernels from regressing widefield activity to stimulus onset times. These kernels correspond to stimulus-driven cortical activity with movement-driven activity removed. Kernels had a temporal component including lags of −300 to 850 ms relative to the stimulus onset. Plots are the maximum between 0-200 ms which are averaged across mice (visuomotor, n = 6 mice; audiomotor, n = 5 mice). Gray outlines represent areas from the Allen CCF. **d**, Cortical stimulus responses as in (c), for well-trained stages, defined as days 4-5 after learning. **e**, Stimulus-responsive areas in the visual and auditory task (permutation test for each pixel comparing the average response 300 ms before stimulus onset and 200 ms after stimulus onset, p<0.05). The medial prefrontal cortex (mPFC) is responsive only to learned visual stimuli, while the anterior prefrontal cortex (aPFC) is responsive to both learned visual and auditory stimuli. **f**, Left, time course of stimulus responses in different brain regions, “R” and “L” denote left and right hemisphere; right, initial activity time defined as the maximum time of the derivative of the fluorescence. Error bars are mean ± s.e. across mice. The aPFC_R_ was active earlier in the auditory tasks than in the visual tasks (shuffle test between visual and auditory tasks: p_aPFC_L_ = 0.055; p_aPFC_R_ = 0.024).

Mice were able to learn the association in either modality. We defined learning by the reaction times being more consistent than expected from chance, with reaction times measured as the latency from stimulus onset to the onset of wheel movement (**Extended Data Fig. 1a**). Consistent reaction times indicated that the mice were turning the wheel in response to stimulus onset, and therefore carrying out a sensorimotor association from stimulus to wheel movement (**Extended Data Fig. 1b-c**). Reaction times converged across days to a stable value in both tasks, and were slightly faster for auditory stimuli compared to visual stimuli, consistent with the faster speeds of the auditory system^10,27^(**Extended Data Fig. 1d**). Both modalities were learned at the same rate, with mice learning either the visual or auditory association in approximately 5 days (**Extended Data Fig. 1e**). We considered days 4-5 after learning (approximately training days 9-10) as the well-trained stage, and performance at this point was comparable between the visuomotor and audiomotor tasks measured as the consistency of reaction times relative to chance (**Extended Data Fig. 1f**).

We used a decoding approach with widefield calcium imaging to isolate stimulus-driven activity across the dorsal cortex. Our goal was to determine how cortical responses to stimuli changed with learning, so we recorded cortical widefield calcium signals throughout training^28^. In this task, sensory-related activity is closely followed and often overlapping with motor-related activity, making it necessary to remove as much of the motor-related component as possible to examine sensory responses (**Extended Data Fig. 1a**). We employed a decoding-based approach to accomplish this, using ridge regression to predict stimulus onsets from cortical activity. The resulting kernels revealed patterns of cortical activity which consistently followed stimulus onset, and importantly did not include activity in the somatomotor cortex, indicating no contamination by movement-related activity (**Extended Data Fig. 2a-b**). This decoding approach was superior to an encoding-based approach of predicting widefield activity from task and behavioral events, in which stimulus kernels were susceptible to contamination by movement-related activity (**Extended Data Fig. 2c**).

The prefrontal cortex developed stimulus responses after learning in modality-specific subregions. Before mice learned the sensorimotor association, visual and auditory stimuli only elicited activity in sensory cortical regions, being the left visual cortex (contralateral to the stimulus) or bilateral auditory cortex (**Fig. 1c**). In mice that learned the visual task, the prefrontal cortex developed a stimulus response as previously described^3^(**Fig. 1d top**). Mice that learned the auditory task also developed a stimulus response in the prefrontal cortex, but notably in a subset of the region activated after visual learning (**Fig. 1d bottom**). When comparing learning-related stimulus responses across modalities, we found that the medial prefrontal cortex (mPFC) was activated unilaterally by only visual stimuli, while the anterior prefrontal cortex (aPFC) was activated bilaterally by either visual or auditory stimuli (**Fig. 1e-f**). This suggests that, while both the mPFC and aPFC develop stimulus responses after learning, the mPFC is specialized for visual stimuli and the aPFC is multimodal.

Stimulus responses followed a temporal progression from posterior to anterior for both modalities, and were temporally asymmetric across hemispheres for visual stimuli. From stimulus onset, activity began in sensory cortex, followed by the left mPFC (contralateral to the stimulus) for visual stimuli, followed by the bilateral aPFC for both stimuli (**Fig. 1f, Extended Data Fig. 3a-b**). While the mPFC exhibited intermediate timing of activation for visual stimuli, the temporal delay from sensory regions to the aPFC was similar between the auditory task and the visual task (**Fig. 1f**). This consistency of aPFC timing across modalities, along with the absence of mPFC activity for auditory stimuli, s uggests that visual activity may not propagate from the mPFC to the aPFC. However, visual responses emerged first in the left aPFC (contralateral to the stimulus) and then in the right aPFC, whereas auditory responses appeared simultaneously in both hemispheres (**Fig. 1f, right**). This suggests that visual responses may propagate across hemispheres from the left to right aPFC.

Learning-induced prefrontal stimulus responses were also present during passive stimulus presentations. To determine whether the observed prefrontal responses were related only to sensory events or modulated by task engagement, we examined cortical activity during passive stimulus presentations. These presentations were given after each training session, and consisted of 50 repeats of 500 ms of each stimulus. Turning the wheel during this phase had no effect, and mice were quiescent on most stimulus presentations. After excluding all trials with wheel movement, prefrontal stimulus responses were observed in the average activity, consistent with the patterns observed in the task stimulus decoding kernels and reinforcing the finding that these responses are not related to movement (**Extended Data Fig. 4a-b**). In order to compare stimulus responses during passive and task contexts, we created decoding kernels for these passive presentations, which closely resembled average activity but removed the effects of ongoing spontaneous activity, most prominently in the retrosplenial cortex (**Extended Data Fig. 3c-d, Extended Data Fig. 4c-d**). Stimulus responses to passive presentations of task stimuli closely matched those observed during task performance, notably including mPFC responses to visual stimuli and aPFC responses to both visual and auditory stimuli (**Fig. 2a**).

**Figure 2.**
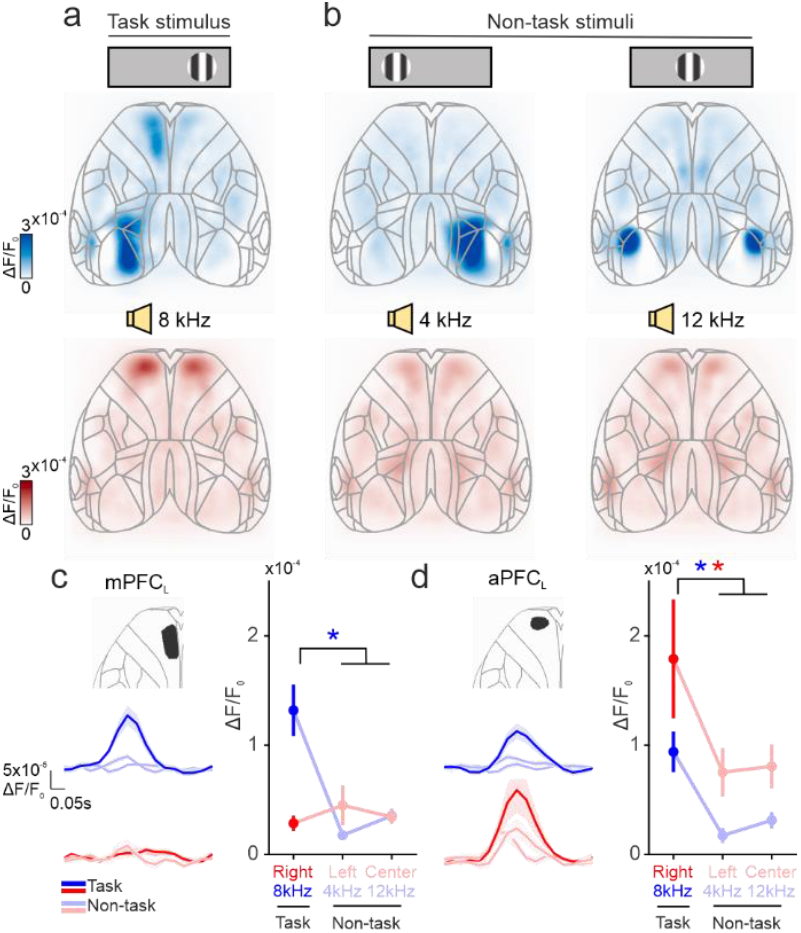
Passive PFC stimulus responses are selective to action-associated stimuli. **aa**, Cortical responses to passively presented stimuli used in the visual (top) or auditory (bottom) task in the well-trained stage. Responses are expressed as kernels predicting stimulus onset times as in Fig. 1c-d. The maximum value between 0–200 ms after stimulus is taken, and maxima are averaged across mice. **b**, Cortical stimulus responses as in (a), for alternate visual (top) or auditory (bottom) stimuli not used during the task. Alternate visual stimuli include a grating on the left or center screen, and alternate auditory stimuli include a 4 kHz and 12 kHz tone. **c**, Left, time course of mPFC responses to the 3 passively presented visual stimuli. Dark colors, task stimulus; light colors, non-task stimuli. Right, maximum mPFC response to passively presented stimuli, error bars are mean ± s.e across mice. The mPFC is selectively responsive to learned visual stimuli (shuffle test between task stimuli and average of 2 non-task stimuli, visual (blue) p = 0.001; auditory (red) p = 0.81). **d**, Plots as in (c) for aPFC responses. The aPFC is selectively responsive to both learned visual and auditory stimuli (shuffle test between task stimuli and average of 2 non-task stimuli, visual (blue) p = 0.003; auditory (red) p = 0.034).

Prefrontal stimulus responses were selective to stimuli trained in the task. During passive stimulus presentation, two additional stimuli were presented for each modality that were not used in the task. For visual stimuli, this included gratings on the center and left screens, and for auditory stimuli, this included a 4 kHz and 12 kHz tone. Cortical responses to non-task stimuli were limited to sensory regions (**Fig. 2b**). The mPFC was selectively active to the trained visual stimulus, and was not active to either non-trained visual stimuli or any auditory stimuli (**Fig. 2c left**). Similarly, the aPFC was more strongly active for trained visual or auditory stimuli compared to non-trained stimuli (**Fig. 2d right**).

Together, these results indicate that learning any sensorimotor association drives the emergence of stimulus responses in the prefrontal cortex, but that the mPFC is specific to visual stimuli while the aPFC is modality-general.

### Context-dependence in prefrontal stimulus responses

Learned stimulus responses in the prefrontal cortex were sensitive to context in a region-specific manner. Visual responses in the mPFC were present in both the task and passive contexts from the first day of visuomotor associative behavior, and increased across days alongside performance (**Fig. 3a**). This resulted in mPFC visual responses that were well-correlated with day-to-day performance of the visuomotor association across contexts (**Fig. 3b**). Conversely, the mPFC never developed a response to learned auditory stimuli, regardless of audiomotor association performance (**Fig. 3c-d**). Visual responses in the mPFC were therefore consistently related to visuomotor performance across contexts, while auditory responses were absent and unrelated to audiomotor performance (**Fig. 3e**). This suggests a modality-specific role of the mPFC in visuomotor learning.

**Figure 3.**
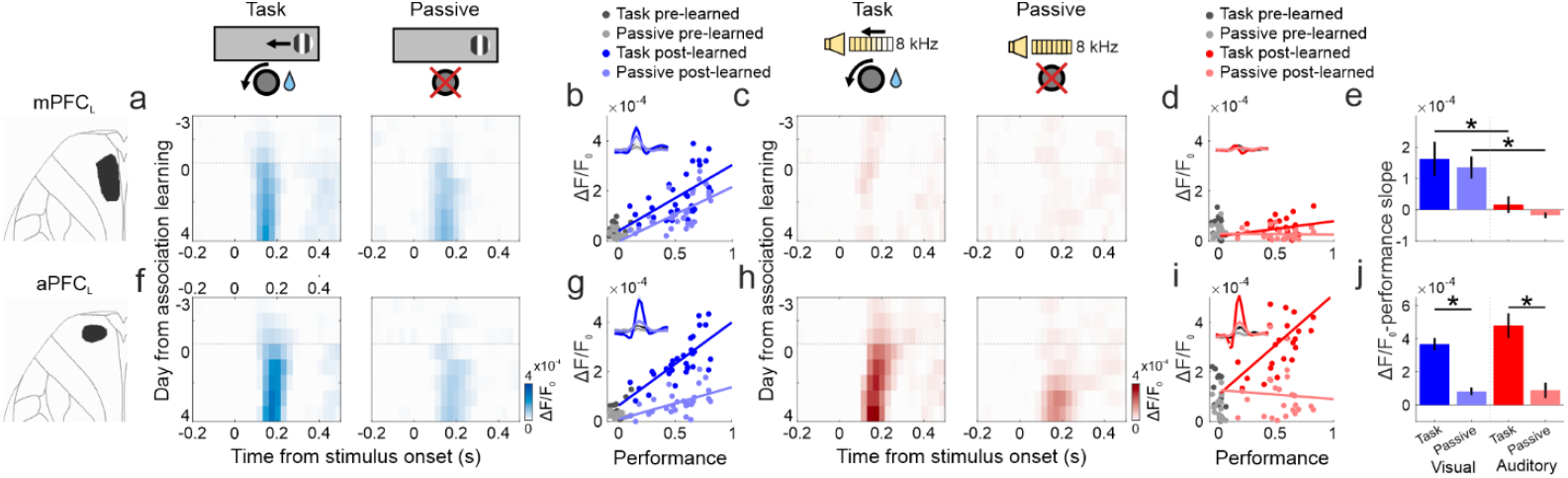
mPFC stimulus responses correlate with performance and are context-independent, while aPFC responses are more variable. **a**, Visual responses in the mPFC across visuomotor learning, aligned to the first day that each mouse exhibited sensorimotor associative behavior (black horizontal line). Left, responses during the task; right, responses during passive stimulus presentation. Responses are expressed as kernels predicting stimulus onset as in Fig. 1-2. **b**, Maximum visual response vs. task performance in the mPFC across visuomotor learning. Task performance is defined as the difference divided by the sum of the reaction time median absolute deviation, such that more consistent reaction times compared to chance yield higher values. Visual responses during the task and passive stimulus presentation are correlated with visuomotor performance (Pearson correlation, r_task_ = 0.59, p_task_ < 0.001; r_passive_ = 0.61, p_passive_ < 0.001), **c**, Responses as in (a) for auditory stimuli across audiomotor learning. **d**, Maximum auditory response vs. task performance in the mPFC as in (b) for audiomotor learning. There is no correlation between audiomotor task performance and auditory responses in the in the mPFC (Pearson correlation, r_task_ = 0.37, p_task_ = 0.07; r_passive_ = 0.05, p_passive_ = 0.82). **e**, Slopes of best-fit lines between stimulus responses and task performance in the mPFC from (b,d). Error bars are mean ± s.e across mice. There is no difference between slopes during the task and passive stimulus presentation in mPFC either during visual and auditory tasks, suggesting a context-independent effect of learning on stimulus responses (shuffle test, visual task vs. passive: p = 0.119; auditory task vs. passive: p = 0.118; task visual vs. auditory: p = 0.005; passive visual vs. auditory: p = 0.002). **f**, Visual responses as in (a), for the aPFC. **g**, Maximum visual response vs. task performance as in (b), for the aPFC. Visual responses in the aPFC are correlated with task performance in both the task and passive contexts (Pearson correlation, r_task_ = 0.75, p_task_ < 0.001; r_passive_ = 0.54, p_passive_ < 0.001). **h**, Auditory responses as in (c), for the aPFC. **i**, Maximum auditory response vs. task performance as in (d), for the aPFC. Auditory responses in the aPFC are correlated with performance only during the task, not during passive stimulus presentation (Pearson correlation, r_task_ = 0.57, p_task_ < 0.001; r_passive_ = 0.06, p_passive_ = 0.76). **j**, Slopes of best-fit lines between stimulus responses and task performance as in (e), for the aPFC from (g,i). Slopes are significantly higher for both modalities during the task compared to passive stimulus presentation, suggesting a context-dependent effect of learning on stimulus responses (shuffle test, visual task vs. passive: p = 0.001; auditory task vs. passive: p = 0.003; task visual vs. auditory: p = 0.905; passive visual vs. auditory: p = 0.797).

In the aPFC, stimulus responses were consistently related to performance only in the task context, and were more variable in the passive context. Like the mPFC, the aPFC exhibited visual stimulus responses in the task from the first day of visuomotor associative behavior (**Fig. 3f**). However, unlike the mPFC, there was an inconsistent relationship between performance and visual responses across the task and passive contexts (**Fig. 3g**). This was also true of audiomotor learning, where auditory responses in the aPFC were correlated to performance only in the task context and not the passive context (**Fig. 3h-i)**. The aPFC therefore exhibited a consistent relationship between stimulus responses and performance only during the task, with passive responses that only weakly increased with improved task performance (**Fig. 3j**). This context dependence in the aPFC did not differ between visual or auditory responses, further suggesting that aPFC stimulus response properties are general across modalities. Together, these results show that stimulus responses in the aPFC are modulated by context, while the mPFC is more purely stimulus-driven regardless of context.

### Cross-modal behavior and prefrontal stimulus responses

Mice were able to transfer visuomotor learning to the audiomotor task, but not the reverse. Having established that mice could learn either the visual or auditory sensorimotor association task, we sought to determine whether mice could transfer learning across modalities. After mice had learned either the visual or auditory task, we then switched the task modality (**Fig. 4a**). Even though the visual and auditory stimuli were initially learned in the same number of days (**Extended Data Fig. 1e**), mice that were switched from visual to auditory stimuli (V_1_A_2_ mice) learned much more quickly than the reverse (A_1_V_2_ mice), with most V_1_A_2_ mice exhibiting associative behavior from the first day of the auditory task (**Fig. 4b, Extended Data Fig. 5a**). The difference in learning transfer between modalities was also apparent in the number of uncued wheel turns when no stimulus was present. Switching modality had no effect on the proportion of uncued wheel turns in V_1_A_2_ mice, while these turns greatly increased after the modality switch in A_1_V_2_ mice, suggesting exploratory behavior due to the lack of a learned cue (**Extended Data Fig. 5b**).

**Figure 4.**
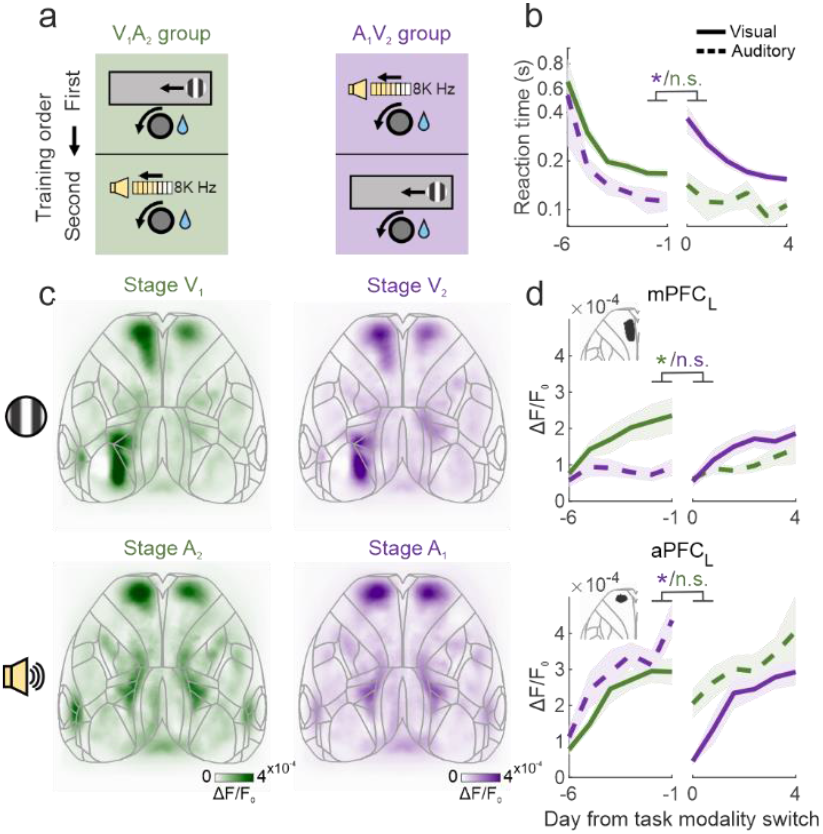
Learning transfers unidirectionally from visual to auditory stimuli, and prefrontal stimulus responses are consistently related to learning. **aa**, Cross-modal learning paradigm. V_1_A_2_ mice learned the visual task followed by the auditory task, and A_1_V_2_ mice learned the auditory and then visual task. **b**, Reaction times for V_1_A_2_ mice (green) and A_1_V_2_ mice (purple) during cross-model learning aligned to first day of the second modality. Solid lines indicate visual task and dotted lines indicate auditory task. Lines and shading are mean ± s.e across mice. A_1_V_2_ mice increased reaction times after switching modalities, while V_1_A_2_ mice displayed consistently low reaction times after switching modalities (shuffle tests on the difference between the average of the last two days of modality 1 and the first two days of modality 2, p_V1A2_ = 0.151, p_A1V2_ = 0.027). **c**, Cortical stimulus responses in the task during well-trained stages of each task modality from V_1_A_2_ mice (left, green) and A_1_V_2_ mice (right, purple). Responses are plotted as in Fig. 1d. The patterns of responses are consistent for learned stimuli, regardless of the order in which they are learned. **d**, Maximum stimulus response in the mPFC (top) and aPFC (bottom) during cross-model learning aligned to first day of the second modality (vertical dotted line). The mPFC response drops after the modality switch in V_1_A_2_ mice (top, green line) despite consistently good performance (b, green line) because it is only responsive to visual stimuli (shuffle test as in (b), mPFC p_V1A2_ = 0.014, p_A1V2_ = 0.977). The aPFC response drops after modality switch only in A_1_V_2_ mice (shuffle test as in (b), aPFC p_V1A2_ = 0.244, p_A1V2_ = 0.027), mirroring the increase in reaction time (b, purple line).

Learned stimulus responses in the prefrontal cortex followed the same regional patterns regardless of learning order. Given the asymmetry in behavioral transfer between modalities, we investigated whether prefrontal stimulus responses were affected by the order in which modalities were learned. For either order, after a given modality was learned, visual stimuli evoked activity in the left mPFC and bilateral aPFC, while auditory stimuli evoked activity only in the bilateral aPFC (**Fig. 4c**). Cortical responses to learned stimuli were therefore not different across groups with different learning orders, indicating that the asymmetry in behavioral transfer was not due to a difference in cortical regions involved.

The dynamics of prefrontal stimulus responses differed by learning order and tracked behavioral associations. Visual responses in the mPFC emerged across visual training, which was the first modality in V_1_A_2_ and the second modality in A_1_V_2_ mice (**Fig. 4d, top solid lines**). This was consistent with visuomotor performance, which required days of training regardless of modality order (**Fig. 4b, Extended Data Fig. 1c**). On the other hand, the modality-general aPFC exhibited an increased response across training in the first modality, regardless of whether it was visual or auditory. After switching the modality, the aPFC response remained high for V_1_A_2_ mice, while it initially dipped and then increased over days for A_1_V_2_ mice (**Fig. 4d bottom**). This mirrored the behavior of each group; where V_1_A_2_ mice immediately performed well on the second modality, A_1_V_2_ mice required days to learn the second modality (**Fig. 4b, Extended Data Fig. 5a**). In summary, the emergence of prefrontal stimulus responses coincided with learning a given modality, which happened at different rates depending on the modality learning order.

Passive prefrontal stimulus responses differed in persistence and amplitude depending on the trained order of modalities. Patterns of cortical responses to passively presented learned stimuli were consistent across either learning order (**Fig. 5a-b**). However, learned visual responses were long-lasting, while auditory responses were transient. For V_1_A_2_ mice which learned the visual association first, both the mPFC and aPFC developed a passive visual response across visuomotor training, both of which persisted after these mice were switched to the audiomotor task (**Fig. 5c top**). In contrast, for A_1_V_2_ mice which learned the auditory association first, the aPFC developed a passive auditory response across audiomotor training, which then went away after visuomotor training (**Fig. 5d bottom right**). The V_1_A_2_ mice also developed a passive auditory response in the aPFC after audiomotor learning (**Fig. 5c bottom right**), but this was much weaker than the response observed in A_1_V_2_ mice (**Fig. 5d bottom right**). Together, these results indicate that learned visual responses in the prefrontal cortex are robust to both context and learning interference, while learned auditory responses are much more sensitive to both factors.

**Figure 5.**
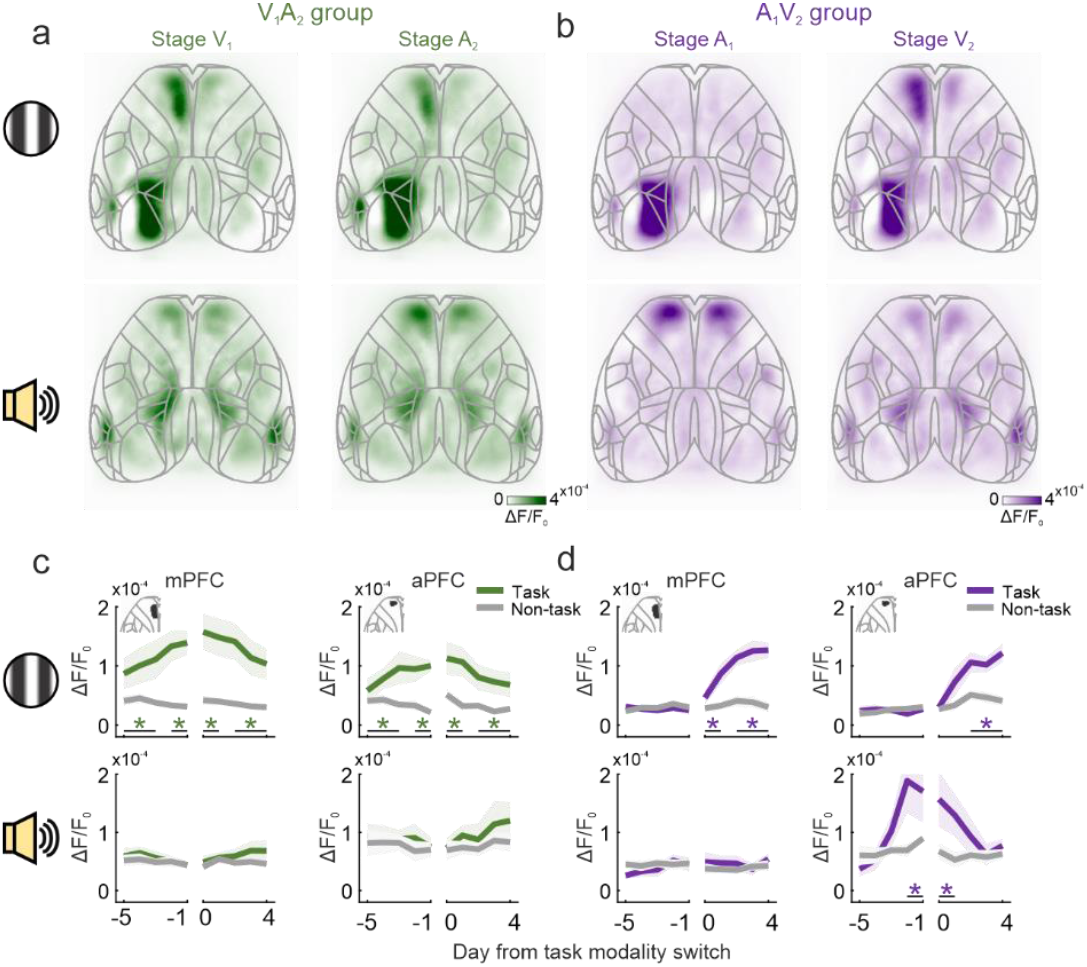
Learning induces a permanent change in mPFC and a transient change in aPFC responses to passively presented stimuli. **aa**, Cortical stimulus responses to passively presented visual (top) and auditory (bottom) stimuli during well-trained stages of the first modality (left) and second modality (right) in V_1_A_2_ mice. Note that the mPFC is still visually responsive after these mice have learned the audiomotor task (top right). **b**, Plots as in (a), for A_1_V_2_ mice. Note that the aPFC is responsive to auditory stimuli after auditory learning (bottom left), and this response is largely absent after switching to the visuomotor task (bottom right). **c**, Maximum stimulus responses in the mPFC (left) and aPFC (right) in V_1_A_2_ mice to the task stimulus (green) and non-task stimuli (gray, average of 2 non-task stimuli) for visual (top) and auditory (bottom) stimuli. Data is aligned to the first day of the second modality (vertical dotted line). Lines and shading are mean ± s.e. across mice. Visual responses in the mPFC and aPFC are consistently larger for task stimuli compared to non-task stimuli across learning, even during audiomotor learning as the second modality (shuffle tests of difference between task and non-task stimuli averaged across 4 groups of days indicated by horizontal black lines, mPFC in visual stimuli: p_day groups_ = 0.007, 0.003, 0.002, 0.002; aPFC in visual stimuli: p_day groups_ = 0.002, 0.004, < 0.001, < 0.001; mPFC in auditory stimuli: p_day groups_ = 0.235, 0.427, 0.267, 0.121; aPFC in auditory stimuli: p_day groups_ = 0.266, 0.396, 0.252, 0.217). **d**, Maximum stimulus responses as in (c), for A_1_V_2_ mice. In A_1_V_2_ mice, the aPFC became responsive selectively to the auditory task stimulus during audiomotor learning, and this response went away across subsequent visuomotor learning (bottom right) (shuffle tests as in (c), mPFC in visual stimuli: p_day groups_ = 0.007, 0.003, 0.002, 0.002; aPFC in visual stimuli: p_day groups_ = 0.316, 0.881, 0.128, 0.004; mPFC in auditory stimuli: p_day groups_ = 0.885, 0.443, 0.239, 0.305; aPFC in auditory stimuli: p_day groups_ = 0.493, 0.029, 0.019, 0.185).

Behavior and cortical responses remained consistent when modalities were mixed in a single task, though kinematics differed by learning order. After mice successfully learned sensorimotor associations for both modalities, they were trained in a task that included both modalities, with either the visual or auditory stimulus being presented at random on each trial (**Extended Data Fig. 6a**). Mice exhibited behavioral associations to both modalities, with similar reaction times, performance, and proportion of uncued wheel movements for each modality regardless of learning order (**Extended Data Fig. 6b-d**). However, the velocity with which mice turned the wheel revealed a behavioral separation between modalities depending on the learning order. Specifically, A_1_V_2_ mice appeared to have two different motor strategies, moving the wheel with high velocity to visual stimuli and with lower velocity to auditory stimuli (**Extended Data Fig. 6e**). This was not just true of the mixed task, but was also observed after learning each modality independently (**Extended Data Fig. 5c**). In contrast, V_1_A_2_ mice appeared to have only one motor strategy, moving the wheel at the same speed regardless of modality. This hints at a relationship between cross-modal transfer and learning strategy, where the V_1_A_2_ mice transferred learning and utilized a single strategy, while the A_1_V_2_ mice did not transfer learning and instead learned two separate strategies. Despite this difference, cortical stimulus responses in the mixed-modality task were similar across groups for both task and passive contexts, indicating that the cortex was not the root of the behavioral difference (**Extended Data Fig. 6f-g**).

### Striatal sensory responses depend on modality learning order

The striatum has an established role in sensorimotor behavior, and is a possible site of difference between mice which learned modalities in different orders. Because mice were able to transfer learned behavior from visual to auditory stimuli, but not the reverse, we hypothesized that the striatum played a role in the transfer and therefore may have different sensory responses based on the modality learning order. This difference would be most likely in an area with overlapping visual and auditory input, which may facilitate learning transfer. The anterior dorsomedial striatum represents such an area, and contains overlapping projections from both the visual and auditory cortex, in contrast to the posterior striatum where these projections are present but segregated (**Fig. 6a-b, Extended Data Fig. 7**).

**Figure 6.**
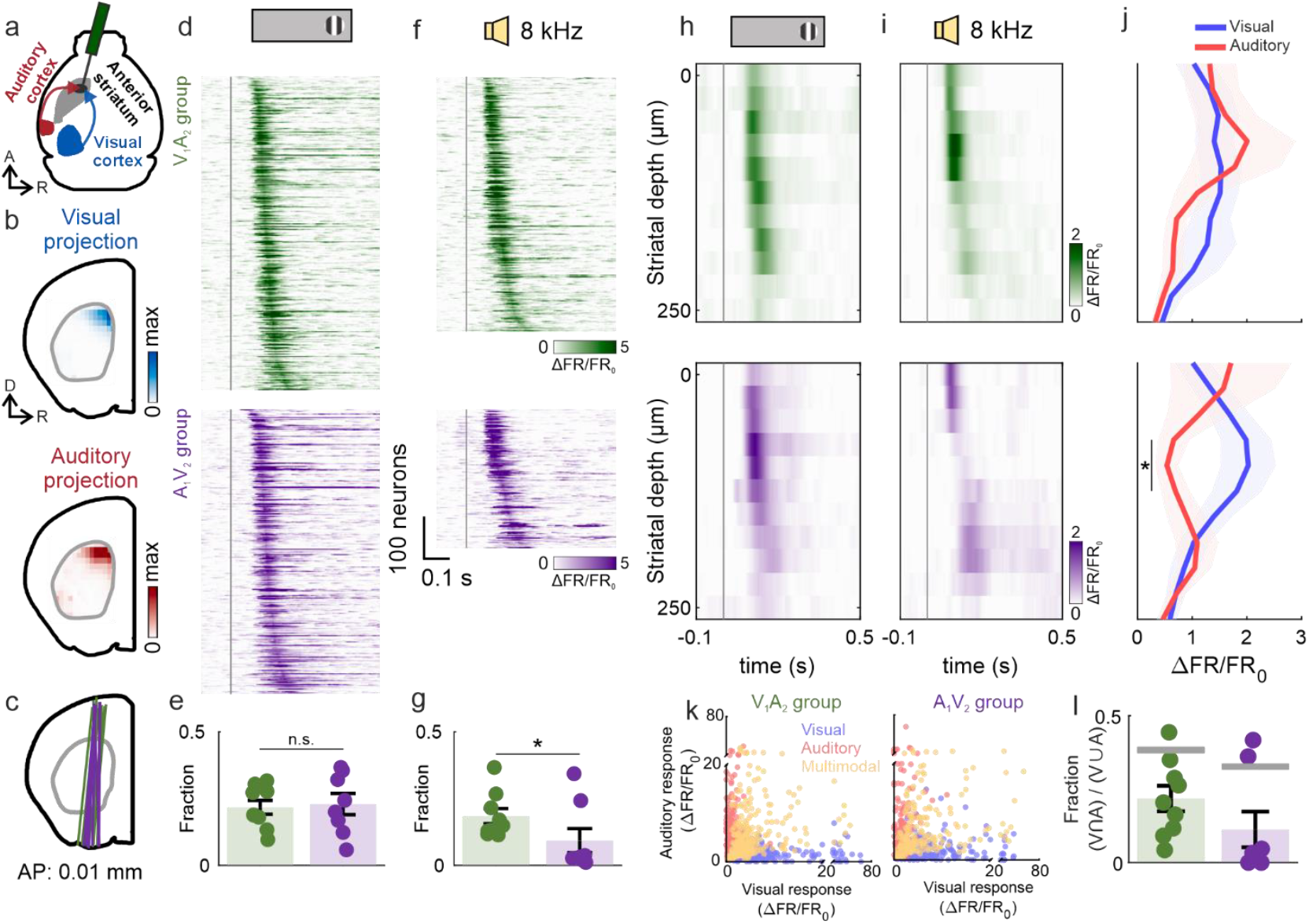
Anterior striatal sensory responses reflect the cross-model learning order. **a**, Schematic of visual and auditory cortical projections to the anterior striatum. **b**, Projection density from visual (top) and auditory (bottom) cortices to the anterior striatum. Data from the Allen Mouse Brain Connectivity Atlas^1^. **c**, Probe trajectory across all recordings (green, V_1_A_2_ mice: 9 recordings across 5 mice; purple, A_1_V_2_ mice: 8 recordings across 4 mice). **d**, Peri-stimulus time histograms (PSTH) of neurons in anterior striatum responsive to visual stimuli in V_1_A_2_ (top, green) and A_1_V_2_ mice (bottom, purple) during passive stimulus presentation, concatenated across all recordings from all mice and sorted by maximum response time. Trials including movement 0-200 ms after stimulus onset are excluded. F.R., firing rate. **e**, Fraction of visually responsive neurons relative to all recorded neurons. No significant difference was observed in the proportion of visually responsive neurons between V_1_A_2_ and A_1_V_2_ mice (shuffle test, p = 0.743). **f**, PSTH of neurons as in (d) for auditory stimuli. **g**, Fraction of responsive neurons as in (e) for auditory stimuli. The proportion of auditory-responsive neurons was significantly higher in V_1_A_2_ mice compared to A_1_V_2_ mice (shuffle test, p = 0.046). **h**, Average multiunit firing rates across all recorded neurons to visual stimuli at different depths in the anterior striatum in V_1_A_2_ (top, green) and A_1_V_2_ mice (bottom, purple). **i**, Average multiunit firing as in (h) for auditory stimuli. **j**, Maximum of the average multiunit firing rates across all recorded neurons to visual stimuli (blue) and auditory stimuli (red) between V_1_A_2_ (top) and A_1_V_2_ (bottom) mice at different depths. Black line with * indicates depths where a shuffle test between visual and auditory multiunit responses p < 0.05. There is no difference in visual responses between V_1_A_2_ and A_1_V_2_ mice, while V_1_A_2_ mice have larger auditory responses at striatal depths 75-125 μm compared to A_1_V_2_ mice. **k**, Scatter plot of striatal neuron responses to visual (x-axis) and auditory (y-axis) stimuli in V_1_A_2_ (left) and A_1_V_2_ mice (right), measured as the maximum PSTH within a 0–200 ms window following stimulus onset. Each dot represents a single neuron; blue, red, and orange dots indicate neurons responsive exclusively to visual, exclusively to auditory, and to both modalities, respectively. **l**, Fraction of multimodally responsive neurons relative to neurons responsive to at least one modality. Dots are recordings, error bars are mean ± s.e. across recordings. Gray rectangles represent null distribution of multimodal neurons, being the 5^th^to 95^th^percentile values from randomly shuffling modality responsiveness across neurons that are responsive to at least one modality. The fraction of multimodally responsive neurons is lower than shuffle for both V_1_A_2_ and A_1_V_2_ mice (p_V1A2_ < 0.001; p_A1V2_ < 0.001), indicating that neurons are preferentially unimodal.

We recorded electrophysiological activity from the dorsomedial striatum while mice performed the mixed-modality task. After mice learned associations for both modalities separately and had been further trained on the mixed-modality task, we performed acute recordings with Neuropixels probes in the dorsomedial striatum during mixed-modality task performance and passive stimulus presentation (**Fig. 6c, Extended Data Fig. 8**).

Mice that switched from the visual to auditory task had more auditory-responsive neurons in the dorsomedial striatum than mice that learned in the reverse order. We classified stimulus responsiveness in each striatal neuron during the passive context using trials without wheel movement. Both V_1_A_2_ and A_1_V_2_ mice exhibited a robust number of visually responsive striatal neurons, with no difference between groups (**Fig. 6d-e**). In contrast, the V_1_A_2_ mice had many more auditory-responsive striatal neurons than A_1_V_2_ mice, with most A_1_V_2_ mice having almost no auditory-responsive units (**Fig. 6f-g**). Auditory responses in the dorsomedial striatum were therefore more prevalent in V_1_A_2_ mice, which transferred learning from visual to auditory modalities, compared to A_1_V_2_ mice, which learned the auditory task with no prior experience.

Striatal visual and auditory responses were present in overlapping spatial distributions. We examined the amplitude of multiunit responses to passively presented visual and auditory stimuli by striatal depth, which provided a spatial distribution of responses that was unbiased by response classification. Multiunit visual responses were located throughout the dorsal-ventral axis of our recording trajectory in both V_1_A_2_ and A_1_V_2_ mice, with a bias towards the dorsal region (**Fig. 6h**). In V_1_A_2_ mice, this region also exhibited auditory responses with a similar amplitude (**Fig. 6i-j top**). In contrast, A_1_V_2_ mice showed much weaker auditory responses compared to visual responses (**Fig. 6i-j bottom**), consistent with the smaller fraction of auditory-responsive neurons in these mice (**Fig. 6g**).

Individual striatal neurons were biased towards being unimodally responsive in mice that exhibited cross-modal learning transfer. V_1_A_2_ mice were able to uniquely transfer learning across modalities, and it could be hypothesized that this was facilitated by multimodal neurons that responded to both stimuli. However, striatal neurons in both groups were more unimodal than expected from chance (**Fig. 6k-l**). This unimodality, together with the consistent fraction of visually responsive neurons across groups (**Fig. 6e**), indicates the additional recruitment of auditory-responsive neurons in V_1_A_2_ mice rather than dynamics in the visually responsive population.

## Conclusion

These results reveal that there are both uni- and multimodal zones of the prefrontal cortex (PFC) that respond to behaviorally relevant stimuli, and that the striatum, but not cortex, exhibits different sensory responses based on the order of learned modalities. Both visuomotor and audiomotor learning induced stimulus responses in the anterior PFC (aPFC) preferentially during task performance, whereas the medial PFC (mPFC) responded exclusively to visual stimuli and did so regardless of context. Mice were able to transfer learned sensorimotor associations from visual to auditory stimuli but not vice versa, revealing an asymmetry in learning transfer across modalities. Despite this behavioral asymmetry, cortical responses to learned stimuli were consistent regardless of learning order. In contrast, this behavioral asymmetry was accompanied by activity differences in the anterior striatum, which was consistently responsive to learned visual stimuli, but only responsive to learned auditory stimuli after transferring from visuomotor learning.

Together, these findings demonstrate that learning can re-route sensory information heterogeneously: some downstream regions (mPFC) are modality-specific, some are modality-general (aPFC), and some (anterior striatum) depend on learning history.

### Regional differences in frontal cortex sensory responses

The differences in context dependence and multimodality between the mPFC and aPFC are possibly due to different sources of input. We found that the mPFC was responsive specifically to visual stimuli and stable across contexts, suggesting that visual information is transmitted directly from visually responsive regions. This input may include direct projections from the visual cortex, although these projections are generally sparse^1^. Instead, visual input into the mPFC may be inherited from a nearby region of the anterior cingulate cortex (ACA) near bregma, which is naively responsive to visual stimuli, and receives input from the visual cortex and projects to the mPFC^1,16,29^. Learning may therefore drive potentiation of these visual inputs into the mPFC, causing the mPFC to stably respond to learned visual stimuli.

In contrast to the mPFC, the aPFC was responsive to both learned visual and auditory stimuli, and was predominantly responsive in the task context. The aPFC receives only sparse visual and auditory input^30^, and instead can receive sensory input indirectly from the cortex-recipient and subcortex-recipient thalamus^31-33^, for example specifically from the mediodorsal and ventromedial thalamic nuclei^34^. Thalamocortical input can be gated by factors like value and attention^35-38^and sensory responses in the thalamus are shaped by sensorimotor learning^39^, which can in turn drive stimulus-associated actions. The aPFC may therefore receive multimodal sensory information primarily while performing sensorimotor associations.

It is worth noting that widefield calcium imaging likely does not report activity of deep areas in the frontal cortex, including the orbitofrontal cortex and the prelimbic and infralimbic areas along the medial wall. Visual and auditory responses have been shown to overlap in these areas^10^, though it is possible that auditory responses are distributed differently than visual responses in these deeper regions.

### Modality salience as a reason for asymmetric transfer

Sensory-responsive territory in the prefrontal cortex was allocated differently depending on the modality, such that the mPFC was specifically responsive to visual stimuli, and there was no region specifically responsive to auditory stimuli. This may relate to visual cues being intrinsically less salient to mice than auditory cues, where visual cues may require a dedicated subregion to direct sensory information to motor circuits. Auditory stimuli, particularly at higher intensities, can elicit robust movements in naïve mice^40,41^, providing an innate sensorimotor substrate for audiomotor learning. By contrast, with few exceptions like looming stimuli^42^, visual stimuli do not elicit reflexive responses and instead require experience-dependent engagement of mPFC to drive behavior^43^.

### Shared or separate pathways as a mechanism of modality learning transfer

We found that activity and behavior differed depending on modality training order, which may indicate that different circuits are recruited based on learning history. Specifically, mice could transfer sensorimotor associations from visual to auditory stimuli, and these mice exhibited both visual and auditory responses in the anterior striatum, along with similar movement kinematics in response to both stimuli. These features suggest that these mice were using the same striatal region to execute sensorimotor associations for both modalities. One possible mechanism for this could be the coincidental potentiation of auditory input during visuomotor training through volume transmission of dopamine^44-46^.

Conversely, animals could not transfer sensorimotor associations from auditory stimuli to visual stimuli, and these mice had visual but not auditory responses in the anterior striatum, and exhibited different movement kinematics in response to stimuli according to modality. This may suggest that one circuit is first established specifically to carry out the audiomotor association, while another one is then established to carry out the visuomotor association. In this case, initial audiomotor association may utilize the posterior striatum, which also receives auditory input and has been shown to exhibit potentiation during audiomotor learning^24,47^. Since visual and auditory inputs do not overlap in the posterior striatum (**Extended Data Fig. 7**)^18,19^, the visual inputs would therefore not be coincidentally potentiated by volumetric dopamine transmission, and visuomotor learning would need to occur as a separate process.

In summary, we found that cortical activity patterns reflect modality-specific differences, whereas the effects of cross-modal learning emerge primarily in subcortical regions. This distinction suggests that these structures play different roles in flexible sensorimotor learning.

## Data and code availability

The datasets generated during the current study are available as downloadable files on OSF at https://osf.io/g7b6r.

The code used to analyze the data are available on GitHub at https://github.com/PetersNeuroLab/PetersLab_papers/tree/main/%2BSong_2025.

## Acknowledgements

We thank Yuliia Shevchuk for collecting part of the Neuropixels data, and Randy Bruno, Haron Avgana and Peter Gorman for proofreading the manuscript and providing valuable comments and suggestions.

## Author contributions

A.J.P. and D.S. conceived and designed the study. D.S. collected and analyzed data. A.J.P. and D.S. wrote the manuscript.

## Methods

### Animals

All experiments were performed in accordance with the UK Animals (Scientific Procedures) Act 1986 under personal and project licenses issued by the Home Office. Animals were transgenic mice (tetO-G6s; Camk2a-tTa^28^) both male and female aged 8 weeks or older. Mice were group-housed when possible and maintained on a 12-h light/dark cycle with ad libitum access to food.

### Surgery

For widefield imaging, mice were implanted with a custom titanium headplate. Surgeries were performed under isoflurane anesthesia, with subcutaneous meloxicam and buprenorphine and administered for analgesia. Animals were positioned in a stereotaxic frame on a heated pad, and the scalp was shaved, disinfected with iodine and alcohol, and removed together with the periosteum to expose the skull. The wound margins were sealed with VetBond (World Precision Instruments), and the titanium headplate was secured over the occipital bone using dental cement (C&B Metabond, Parkell). The headplate had a U-shaped component that was centered over the skull, which was filled first with clear dental cement and then UV-curable optical adhesive (NOA81, Norland Products). Meloxicam was provided in the drinking water for 3 days postoperatively.

For electrophysiological recordings, the anterior striatum of the left hemisphere was targeted (approximately 0.5 mm AP and 1.0 mm ML relative to bregma). Mice were anesthetized with isoflurane and given a subcutaneous injection of meloxicam, and were head-fixed via the implanted headplate. The optical adhesive above the target site was drilled away, and a small craniotomy was made with a drill. The craniotomy was covered with Kwik-Cast (World Precision Instruments), and recordings commenced at least 1 day after the procedure.

### Behavioral training

During behavioral training, mice were given 1–5% citric acid water in their home cage to maintain a 10–15% reduction in body weight^48^. Mice not undergoing training had free access to regular water. During behavioral training and recordings, mice were head-fixed in the center of three screens and with their forepaws on a wheel that could be turned leftwards or rightwards^49,50^.

The behavioral task was adapted from Peters et al.^3^with small modifications and implemented in Bonsai^51^. Each trial began with two independent and randomly selected delay periods. The first was an inter-trial interval (ITI) of fixed duration, and the second was a quiescence period during which wheel movements reset the timer until the mice remained still for the required duration. During the first 2 days of training, the ITI ranged from 1–3 s and the quiescence period from 0.5–1 s, both randomized in 100-ms increments; from day 3, these were extended to 4–7 s and 0.5–2 s respectively. After the mouse held the wheel still throughout the quiescence period, the stimulus was presented.

In the visual task, the stimulus was a circle containing a static vertical grating that was displayed on the center of the right-hand screen. The horizontal position of the stimulus was controlled by the wheel, such that clockwise wheel rotations moved it rightward and counterclockwise wheel rotations moved the stimulus leftward.

In the auditory task, the stimulus was 8 kHz pure tone at 470 units of sound intensity (70 dB). Wheel rotations modulated the volume, such that counterclockwise rotations decreased the volume, while clockwise rotations increased the volume.

In both cases, turning the wheel leftwards by 30° would be rewarded with 6 μl of 10% sucrose water, while turning the wheel rightwards by 15° would end the trial. In the visual task, trials were rewarded when the stimulus reached the center, and ended unrewarded when the stimulus went off the right-hand screen. In the auditory task, trials were rewarded when the stimulus was nearly inaudible, and ended unrewarded when the stimulus became louder.

In the mixed-modality task, visual and auditory trials were randomly interleaved throughout each session within a single day, with an equal probability of occurrence. The behavioral contingencies, including wheel rotation thresholds for success or failure and reward delivery, were identical to those described for unimodal tasks.

### Visual and auditory passive stimulus presentation

Visual and auditory stimuli were passively presented for three consecutive days prior to the onset of training, serving two purposes: (1) acclimating the animals to the recording rig, and (2) providing a baseline measurement of cortical responses to visual and auditory stimuli. In addition, the same passive stimulus presentations were also conducted after each day’s task training to evaluate how stimulus-evoked responses were modulated by learning. During passive stimulus presentations, turning the wheel had no effect.

For passively presented visual stimuli, the stimuli consisted of circles containing gratings identical in size and spatial frequency to those used in the task. Stimuli were presented randomly on the left, center, or right screen, with each condition repeated 50 times. Each stimulus was displayed for 500 ms, followed by an inter-stimulus interval randomly selected between 2–3 s in 100 ms increments.

For passively presented auditory stimuli, pure tones at 4 kHz, 8 kHz, and 12 kHz were presented in a randomized order, with each tone repeated 50 times. Tone duration and inter-stimulus intervals matched those of the passive visual stimulus presentation.

The visual stimuli presented on the right-hand screen and the auditory stimuli presented at 8 kHz were identical to those used during training, thereby facilitating direct comparison of cortical activity between task and passive conditions.

### Behavioral performance and statistical definition of learning

A conditional randomization test was applied to evaluate whether mice were moving the wheel in response to the stimulus and therefore had learned the sensorimotor association^3,52^.

Specifically, we tested whether mice initiated the rewarded wheel turn at a consistent time relative to stimulus onset. We did this by quantifying the time from stimulus onset to the onset of the rewarded wheel turn as the reaction time, and comparing the median absolute deviation (m.a.d.) of the reaction time compared to a null distribution.

The null distribution of reaction times was created by sampling from all stimulus onset times that would have been possible given the task parameters and mouse behavior. The actual stimulus onset times were determined by the randomly drawn quiescence times and the wheel turns that reset that quiescence timer. The stimulus onsets therefore fell within a quiescence period between the last timer-resetting movement and the first post-stimulus movement. For each trial, there were a range of other possible quiescence times that would have resulted in stimulus onset during this same quiescence interval, but at a slightly different time. If the mouse was ignoring the stimulus, then the reaction time statistics would be the same using actual stimulus onset times or sampling from all valid stimulus onset times. If the mouse was moving in response to the stimulus onset, then the reaction time statistics should differ when comparing actual to all valid stimulus onset times.

We randomly sampled valid stimulus onset times from each trial, calculated the reaction time relative to the sampled stimulus onsets, and took the m.a.d. of reaction times across trials. We repeated this process 10,000 times to generate a null distribution of reaction time m.a.d. When the measured m.a.d. of reaction times was at p < 0.05 compared to the null distribution, we considered the reaction times more stimulus-locked than chance. In the auditory task, pure tone stimuli occasionally elicited a startle reflex, leading mice to make small, non-task-related wheel movements. To avoid including such artifacts, we defined the onset of the task-mediated response as the beginning of the final continuous wheel rotation occurring before stimulus offset.

To quantify the degree to which mice were consistently responding to the stimulus, we defined “performance” (e.g. **Fig. 3b** x-axis) relative to the measured and null reaction time (RT) m.a.d.:

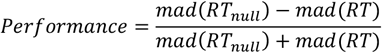

### Widefield imaging and fluorescence processing

Widefield imaging was performed using a Hamamatsu ORCA-Flash4.0 V3 CMOS camera mounted on a macroscope (Scimedia THT-FLSP) with a 1.0x condenser and 0.63x objective (Leica). Images were acquired at 70 Hz with 2×2 binning using a custom MATLAB program to interface with the camera API (DCAM-SDK4). Illumination was produced by a Cairn OptoLED with alternating blue (470 nm, ET470/40x) and violet (405 nm, ET405/20x) excitation, yielding 35 Hz per channel. Blue excitation yielded GCaMP-dependent signals, while violet excitation yielded an activity-independent hemodynamic reference. Excitation light was delivered through the objective via liquid light guide and dichroic, and emission was filtered (525/50–55) before detection.

Widefield data were compressed using singular value decomposition (SVD) of the pixel × time fluorescence matrix, retaining the top 2000 components. All subsequent orthogonally invariant operations (e.g., averaging, deconvolution) were performed on the reduced representation to improve computational efficiency.

Hemodynamic artifacts were corrected by first calculating a scaling factor from the violet signal to the blue signal after spatial downsampling, temporal alignment of alternating frames, and band-pass filtering at roughly the heartbeat frequency (8–12 Hz). The raw violet signal was then linearly detrended, scaled, and subtracted from the blue signal. The resulting hemodynamically corrected signal was normalized by ΔF/(F_0_ +S), with F_0_ baseline defined as the session mean per pixel, and S being a softening factor of the median fluorescence value across all pixels. Finally, the fluorescence was deconvolved with a kernel derived from simultaneous widefield imaging and electrophysiology to best match multiunit spiking^2^.

Experiment-specific SVD components were projected onto a master spatial basis (U_master_) derived from multiple mice, which allowed for aligning temporal components to a single spatial component basis set. This was done by:

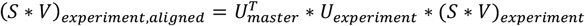

This enabled combining fluorescence data across experiments using a common spatial basis set, reducing computation and memory requirements.

Widefield images were spatially aligned both within and across mice. To align days within mice, the average image from each day was aligned by rigid transformation to the first day of recording. To align across mice, we first created retinotopic maps of visual field sign using sparse noise mapping, which highlighted visual regions^53^. We then aligned the visual field sign map for each mouse to a master visual field sign map by similarity transformation.

Cortical areas from the Allen Mouse Brain Common Coordinate Framework (CCF v3, Allen Institute for Brain Science)^54^were overlaid on widefield images by first assigning values to CCF visual regions based on their expected visual field sign^53^, then aligning that onto the master visual field sign map by similarity transformation.

### Regression between task events and fluorescence

To determine the cortical fluorescence patterns that accompanied stimulus onsets, we performed decoding regression from deconvolved widefield fluorescence to stimulus onset using ridge regression (λ = 15) (e.g. **Fig. 1c**). Widefield signals (N × T) served as regressors, and stimulus onset was modeled as the output, which was expanded with temporal shifts from –0.3 to +0.9 s (–10 to +30 frames at 35 Hz). This produced decoding kernels that capture the temporal structure by which distributed cortical activity predicts the timing of task stimuli. This approach directly tests the predictive information about task events contained in cortical activity, and therefore is not sensitive to extraneous activity like that elicited by movement.

As a comparison to the decoding approach above, we performed encoding regression from task events to deconvolved widefield fluorescence using ridge regression^2^(**Extended Data Fig. 2c**). This is similar to the decoding regression, but predicts fluorescence from task events, rather than predicting task events from fluorescence. Three events were used as regressors: stimulus onset, wheel movement onset, and outcome onset. Each event was represented as a binary time series and expanded with temporal shifts from –300 to +850 ms relative to event onset (–10 to +30 frames at 35 Hz), yielding event-specific design matrices. The regression estimated temporal kernels (regressor × time lag) that describe the event-locked dynamics of cortical activity. These kernels can be strongly affected by the richness of the task regressors; for example, movement-related activity can bleed into stimulus kernels if the movement is not adequately modeled.

### Electrophysiological recordings

Neural recordings were obtained using Neuropixels 3A and 1.0 probes (Jun et al., 2017) mounted on motorized micromanipulators (New Scale Technologies). Each mouse was acutely recorded across multiple sessions. To enable reconstruction of probe trajectories, probes were coated with the fluorescent dye DiI (ThermoFisher) by repeated dipping (5–6 times with brief air drying between dips). Data were acquired using Open Ephys^55^, spike-sorted with Kilosort4^56^, and units representing noise were removed with Bombcell^57^Probe trajectories were reconstructed from histology by aligning coronal sections to the Allen CCF and manually tracing the dye track using publicly available code (https://github.com/petersaj/AP_histology). Because probe endpoints could not be reliably determined from histology and recording depth varied across sessions, electrode depth was refined using electrophysiological landmarks. In particular, the corpus callosum forms a unit-sparse zone between the cortex and striatum. We manually identified the cortical–striatal border as the depth at which the fewest units were detected, typically located in the upper one-third of the probe track. This landmark was used to define the superficial surface of the striatum.

### Cortico-striatal projection maps

Projection data were obtained from the Allen Mouse Brain Connectivity Atlas (http://connectivity.brain-map.org/) to map projections from the visual and auditory cortex to the anterior striatum (**Fig. 6b**, and **Extended Data Fig. 7**). We used Brain Street View (https://github.com/Julie-Fabre/brain_street_view) to access this data and visualize projection density from the visual and auditory cortex to the anterior striatum.

## Extended Data Figures

**Extended Data Figure 1.**
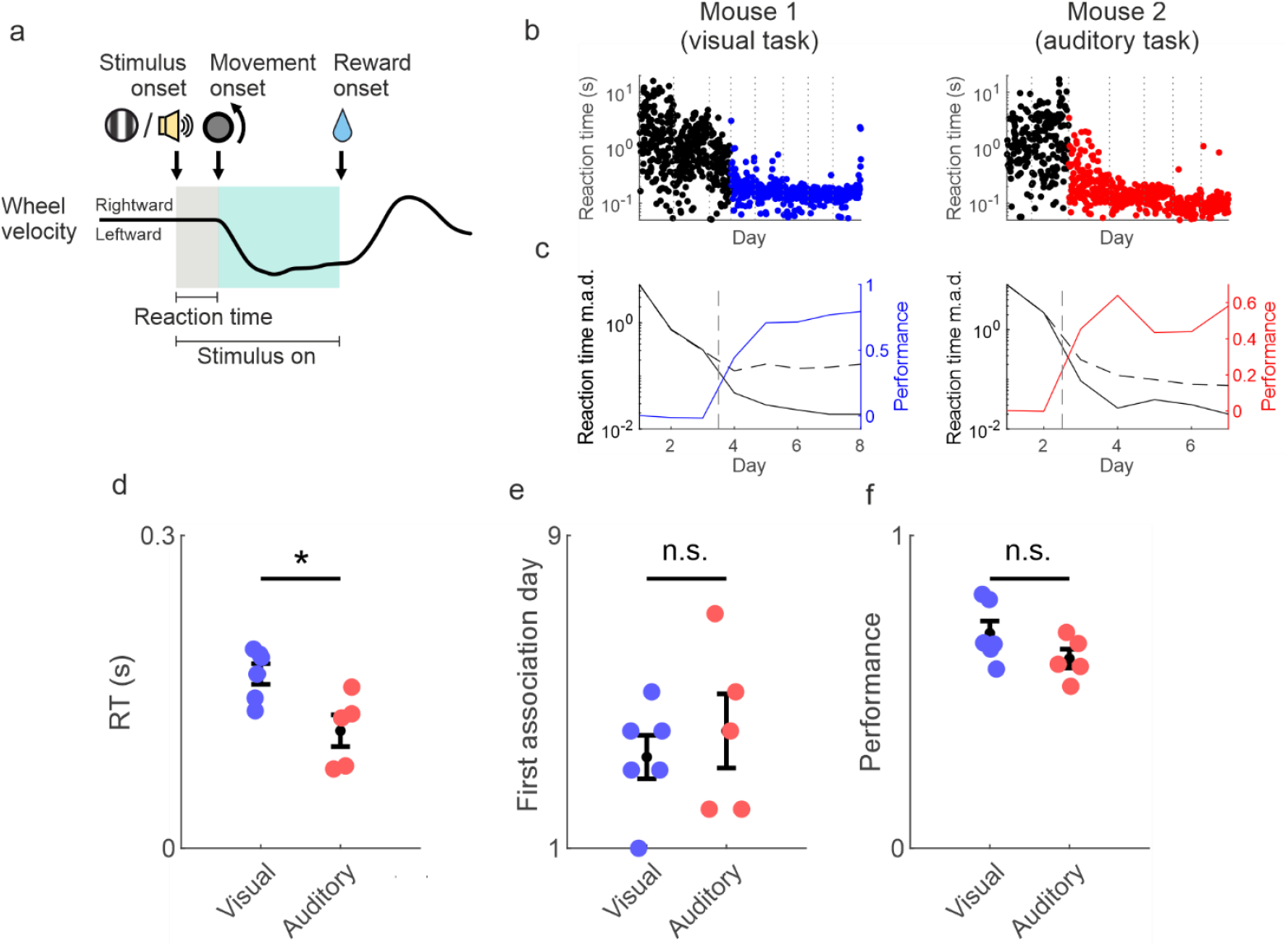
Behavioral performance for visuomotor and audiomotor tasks. **a**, Events in an example trial. **b**, Examples from two mice of reaction time on each trial across days in visual task (left) and auditory task (right). Each dot represents a trial, vertical dotted lines denote days. Black dots indicate trials on days before mice learned the task, colored dots (blue, visual task; red, auditory task) indicate trials after mice had reached the learning criterion. **c**, Median absolute deviation (m.a.d., black lines) and null m.a.d. expected from randomly timed movements (dotted lines) plotted on the left y-axis, and performance (colored lines) as the difference divided by the sum of the measured and null reaction time m.a.d. plotted on the right y-axis, across days in the visual (left) and auditory (right) tasks. Vertical gray line denotes first day where the reaction time m.a.d. was significantly smaller than the null, which represented the first day that mice exhibited sensorimotor association behavior. **d**, Average reaction time for the visuomotor (blue) and audiomotor (red) tasks during the well-trained stage, defined as days 4-5 after learning. Dots are mice, error bars are mean ± s.e. across mice. Reaction times for auditory stimuli are faster than for visual stimuli (difference compared to shuffling modalities, p = 0.016). **e**, Number of days until sensorimotor associative behavior was first exhibited for the visuomotor (blue) and audiomotor (red) tasks. Dots are mice, error bars are mean ± s.e. across mice. There is no difference by modality (difference compared to shuffling modalities, p = 0.713). **f**, Performance, defined as the difference divided by the sum of the measured and null reaction time median absolute deviations, for the visuomotor (blue) and audiomotor (red) tasks during the well-trained stage. Dots are mice, error bars are mean ± s.e. across mice. There is no difference by modality (difference compared to shuffling modalities, p = 0.067).

**Extended Data Figure 2.**
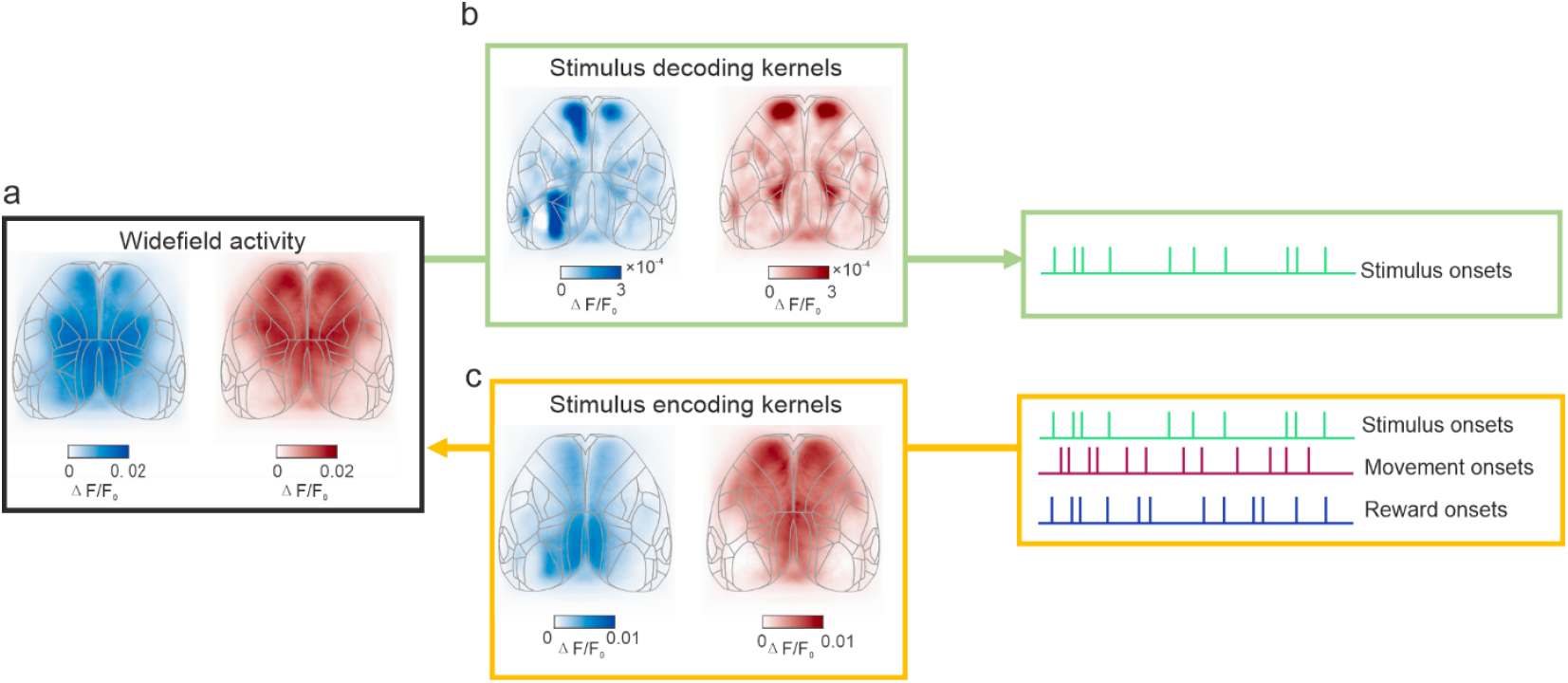
Schematic of calculating decoding kernels and encoding kernels to extract stimulus responses. **a**, Maximum activity 0-200 ms after visual stimulus onset (blue) or auditory stimulus onset (red) during the task, averaged across days in the well-trained stage and mice. **b**, Schematic illustrating decoding regression. Ridge regression is used to create a kernel that estimates stimulus onsets from cortical fluorescence. Kernels have a temporal component that includes lags from −300 to 850 ms, and the maximum from 0-200 ms is shown. Kernels are averaged across well-trained days and mice, corresponding to the images in (a). **c**, Schematic illustrating encoding regression. Linear regression is used to create kernels that estimate total cortical fluorescence from task events including stimulus onsets, movement onsets, and reward onsets. All event kernels have temporal components that include lags from −300 to 850 ms. The stimulus kernel is shown as the maximum value for lags 0-200 ms, and averaged across well-trained days and mice corresponding to the images in (a).

**Extended Data Figure 3.**
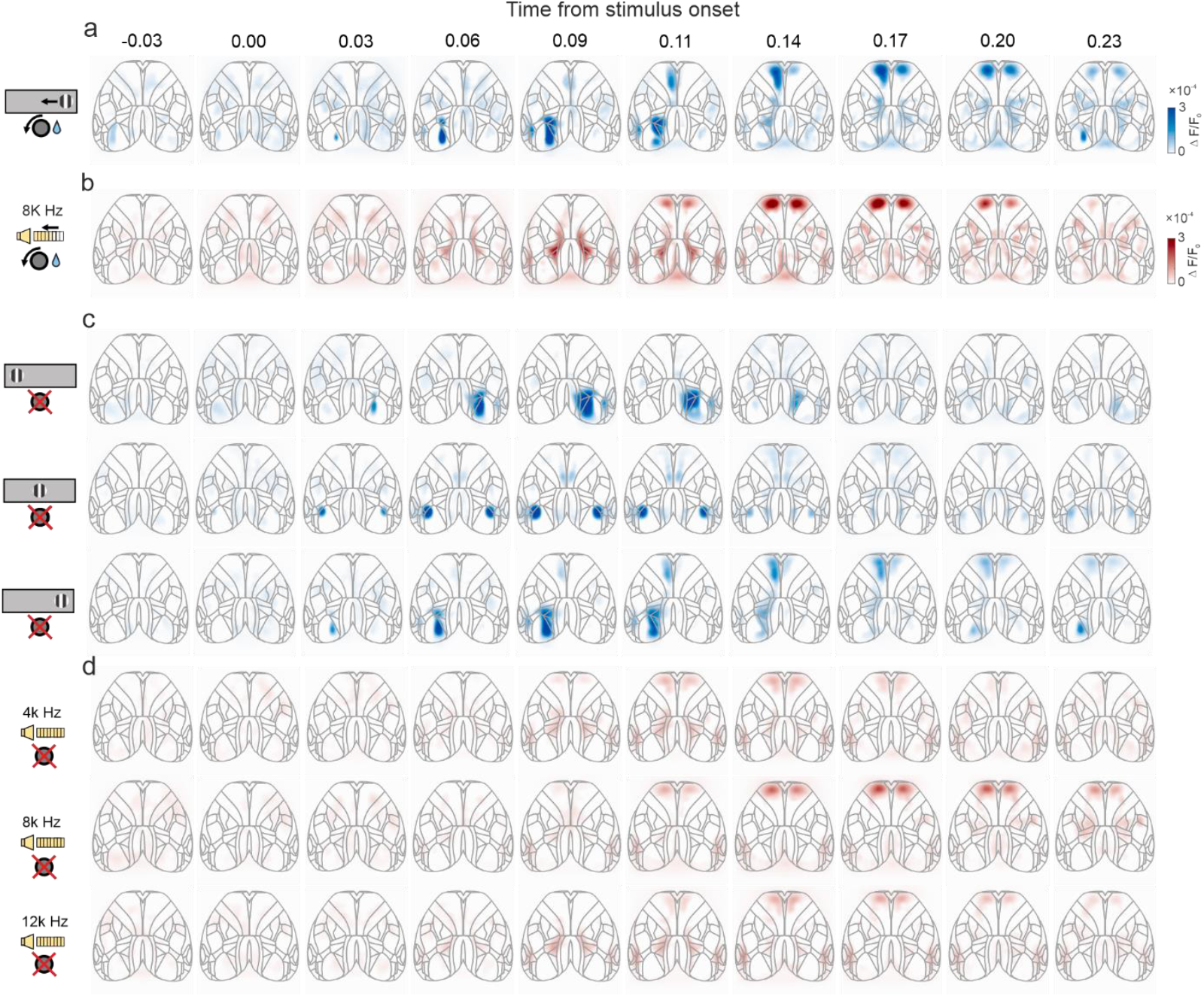
Cortical activity aligned to stimulus onset by timepoint. Average cortical stimulus responses in the well-trained stage by time point relative to stimulus onset in the visual task (a), auditory task (b), visual stimulus passive presentation (c) and auditory stimulus passive presentation (d).

**Extended Data Figure 4.**
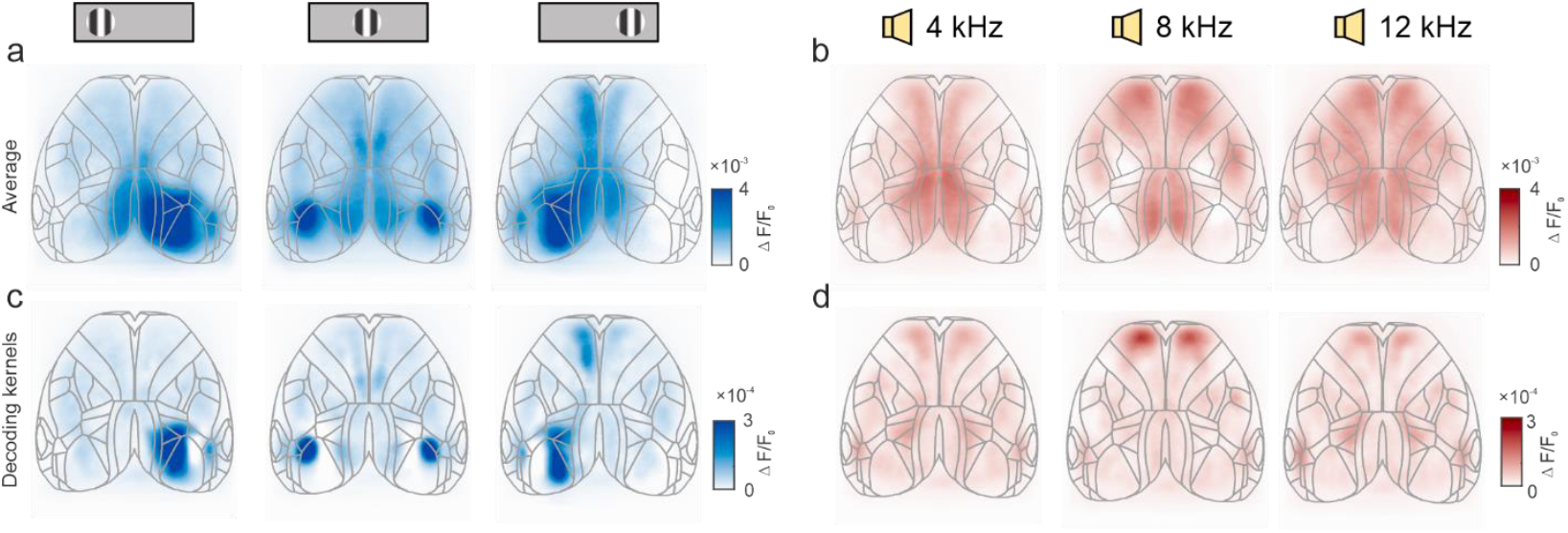
Comparison between total fluorescence and stimulus onset decoding kernels. **a**, Maximum fluorescence 0– 200 ms after visual stimulus onset averaged across well-trained days and mice. Trials with wheel movement during this time were excluded. Columns are responses to visual stimuli on the left, center, and right screens. **b**, Maximum fluorescence responses 0-200 ms after auditory stimulus onset averaged across well-trained days and mice. Columns are responses to auditory stimuli at frequencies of 4 kHz, 8 kHz, and 12 kHz. **c**, Cortical responses to passively presented visual stimuli expressed as decoding kernels. The maximum value from 0-200 ms after stimulus onset is used, then averaged across well-trained days and mice. Columns are responses to visual stimuli on the left, center, and right screens as in (a). These plots are also shown in Fig 2a-b, and are included here for comparison with (a). **d**, Cortical responses to passively presented auditory stimuli expressed as decoding kernels. The maximum value from 0-200 ms after stimulus onset is used, then averaged across well-trained days and mice. Columns are responses to auditory stimuli at frequencies of 4 kHz, 8kHz, and 12 kHz. These plots are also shown in Fig 2a-b, and are included here for comparison with (b).

**Extended Data Figure 5.**
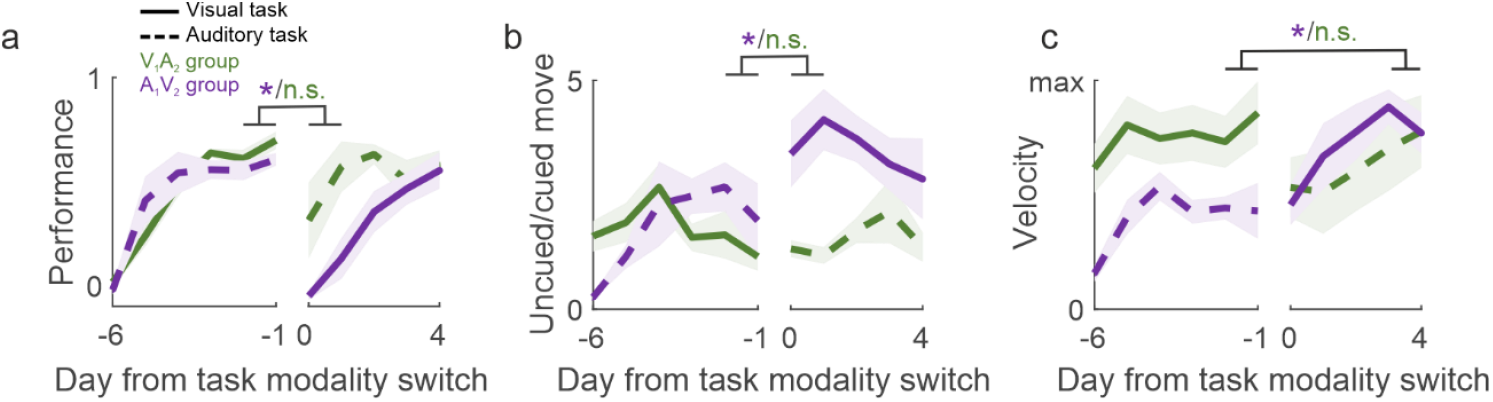
Behavioral measures for V_1_A_2_ and A_1_V_2_ mice across training. **a**, Task performance for V_1_A_2_ mice (green) and A_1_V_2_ mice (purple) during cross-model learning aligned to first day of the second modality. Solid line indicates visual task and dotted line indicates auditory task. Lines and shading are mean ± s.e. across mice. A_1_V_2_ mice reduced performance after switching modalities, while V_1_A_2_ mice displayed consistently high performance after switching modalities (shuffle tests on the difference between the average of the last two days of modality 1 and the first two days of modality 2: p_V1A2_ = 0.221, p_A1V2_ = 0.035). **b**, Number of movements when a stimulus was not present (uncued) divided by the number of movements while a stimulus was present (cued). A_1_V_2_ mice increased their relative number of uncued movements after switching modalities, while V_1_A_2_ mice displayed a consistent ratio of cued and uncued movements after switching modalities (shuffle tests as in (a): p_V1A2_ = 0.594, p_A1V2_ = 0.029). **c**, Trial-averaged maximum wheel velocity in response to the stimulus. In the well-trained stage, A_1_V_2_ mice exhibited higher velocity wheel movements in the visual task than in the auditory task, while V_1_A_2_ mice exhibit similar wheel velocity between the visual and auditory task. (shuffle tests on the difference between the average of the last two days of modality 1 and the day 4 and 5 of modality 2: p_V1A2_ = 0.018, p_A1V2_ = 0.463). Data is aligned to the first day of the second modality (vertical line), lines and shading are mean ± s.e. Note that wheel velocity is separate from reaction time; for example, a mouse can have a fast reaction time and a slow wheel velocity.

**Extended Data Figure 6.**
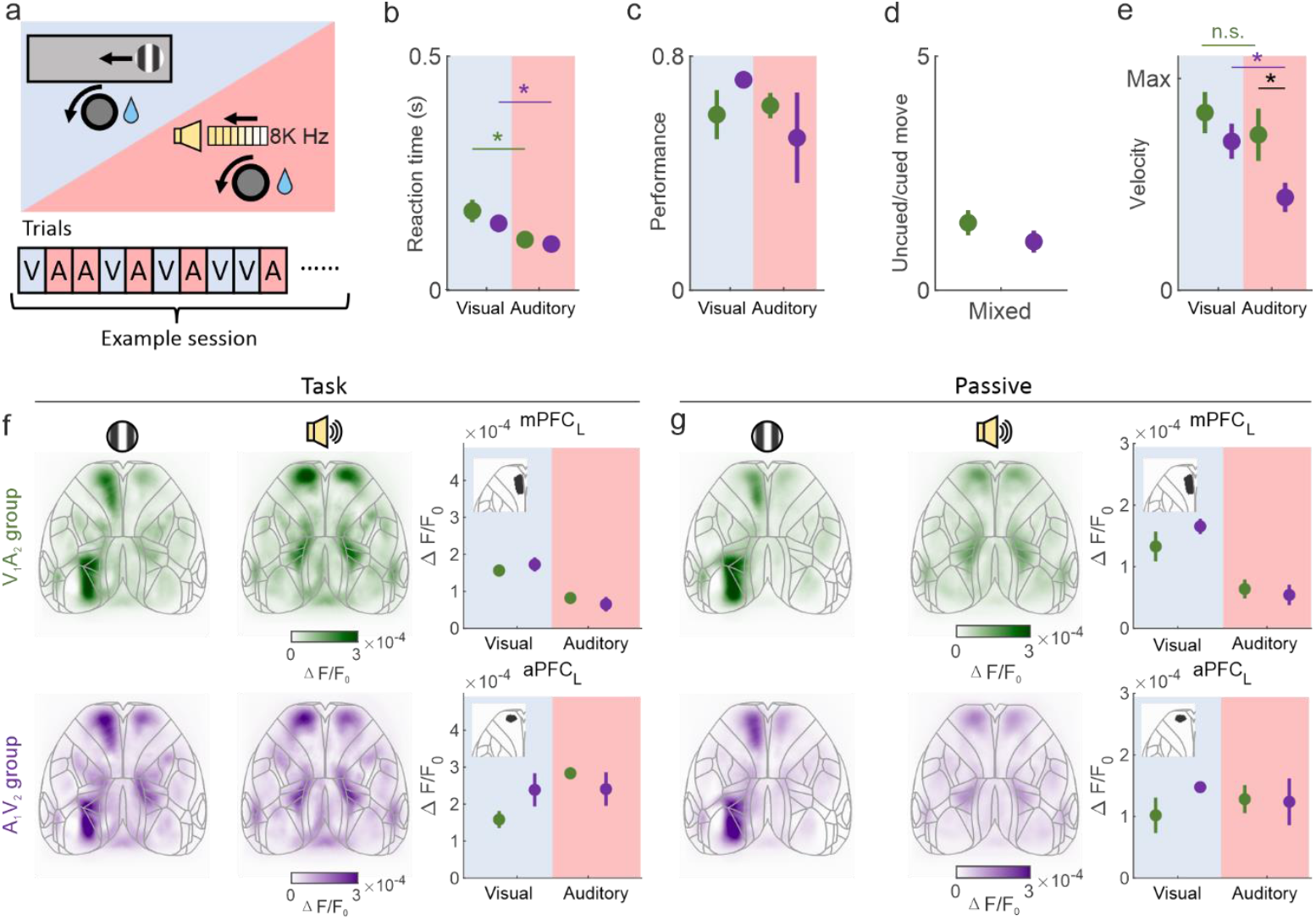
Behavioral and cortical activity in the mixed-modality task. **a**, Behavioral paradigm of mixed-modality task. **b**, Reaction time for V_1_A_2_ (green) and A_1_V_2_ (purple) mice during the mixed-modality task. There is no difference between groups within a single modality, and reaction time in auditory trials is lower than in visual trials in both groups (shuffle test, p_visual_ = 0.353; p_auditory_ = 0.658; p_V1A2_ = 0.018; p_A1V2_ = 0.029). **c**, Performance defined as the difference divided by the sum of measured and null reaction time m.a.d. for V_1_A_2_ and A_1_V_2_ mice during the mixed-modality task. There is no difference between visual and auditory trials within animal groups or between mice groups within single modality (shuffle test, p_visual_ = 0.837; p_auditory_ = 0.318; p_V1A2_ = 0.603; p_A1V2_ = 0.184). **d**, Number of uncued movements (without a stimulus present) divided by the number of cued movements (while a stimulus was present) for V_1_A_2_ and A_1_V_2_ mice during the mixed-modality task. There is no difference between mice groups (shuffle test, p = 0.161). **e**, Trial-averaged maximum wheel velocity in response to the stimulus for V_1_A_2_ and A_1_V_2_ mice. V_1_A_2_ mice turn the wheel with higher velocity than A_1_V_2_ mice during the auditory trials in the mixed task (shuffle test, p_visual_ = 0.127; p_auditory_ = 0.018; p_V1A2_ = 0.285; p_A1V2_ = 0.029). **f**, Left, cortical responses to visual and auditory stimuli during the mixed-modality task for V_1_A_2_ and A_1_V_2_ mice; right, maximum responses in the mPFC (top) and aPFC (bottom), error bars are mean ± s.e. across mice. There is no difference in cortical responses between V_1_A_2_ and A_1_V_2_ group in mPFC and aPFC during visual and auditory trials in mixed tasks (p_mPFC-visual_ = 0.349; p_mPFC-auditory_ = 0.505; p_aPFC-visual_ = 0.836; p_aPFC-auditory_ = 0.558). **g**, Left, cortical responses to passively presented visual (left) and auditory (right) stimuli for V_1_A_2_ and A_1_V_2_ mice; right, maximum responses in the mPFC (top) and aPFC (bottom) for passively presented task stimuli during days where mice performed the mixed-modality task. Error bars are mean ± s.e. across mice. There is no difference in mPFC or aPFC responses between groups (p_mPFC-visual_ = 0.733; p_mPFC-auditory_ = 0.332; p_aPFC-visual_ = 0.854; p_aPFC-auditory_ = 0.119).

**Extended Data Figure 7.**
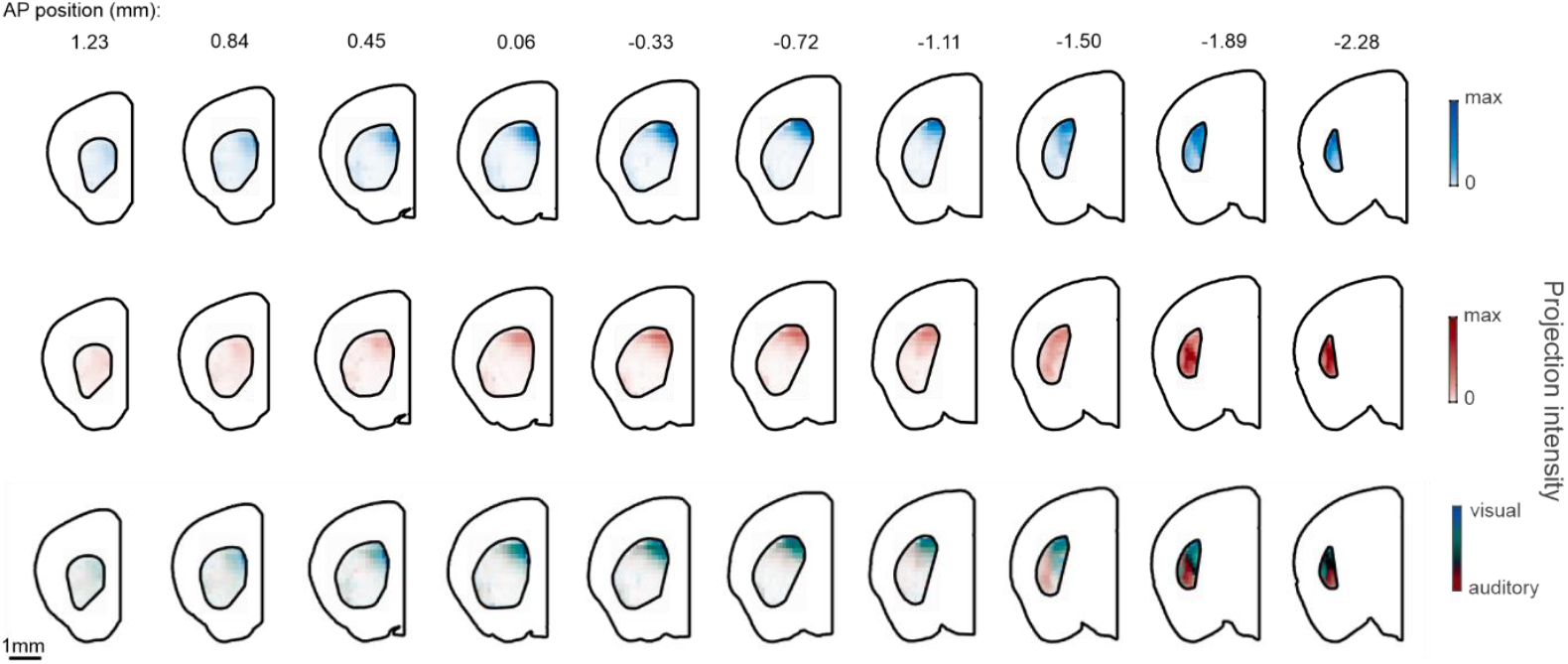
Distribution of visual and auditory cortical inputs along the striatal anterior–posterior axis. Projection distributions from the auditory and visual cortices to the striatum (anterior-to-posterior as left-to-right) obtained from the Allen Mouse Brain Connectivity Atlas^1^. Slices are spaced 0.39 mm apart, and the values above each panel indicate the distance anterior or posterior to bregma. The heatmap shows the density of axonal projections from the visual (blue) or auditory (red) cortex at each position, with darker colors indicating higher projection intensity. Values are normalized to the maximum pixel value across all slices. The bottom merged row shows the overlay of auditory (red) and visual (blue) projections in each slice, illustrating regions of overlap or segregation along the anterior–posterior axis. Note that visual and auditory inputs overlap in the anterior striatum and are segregated in the posterior striatum.

**Extended Data Figure 8.**
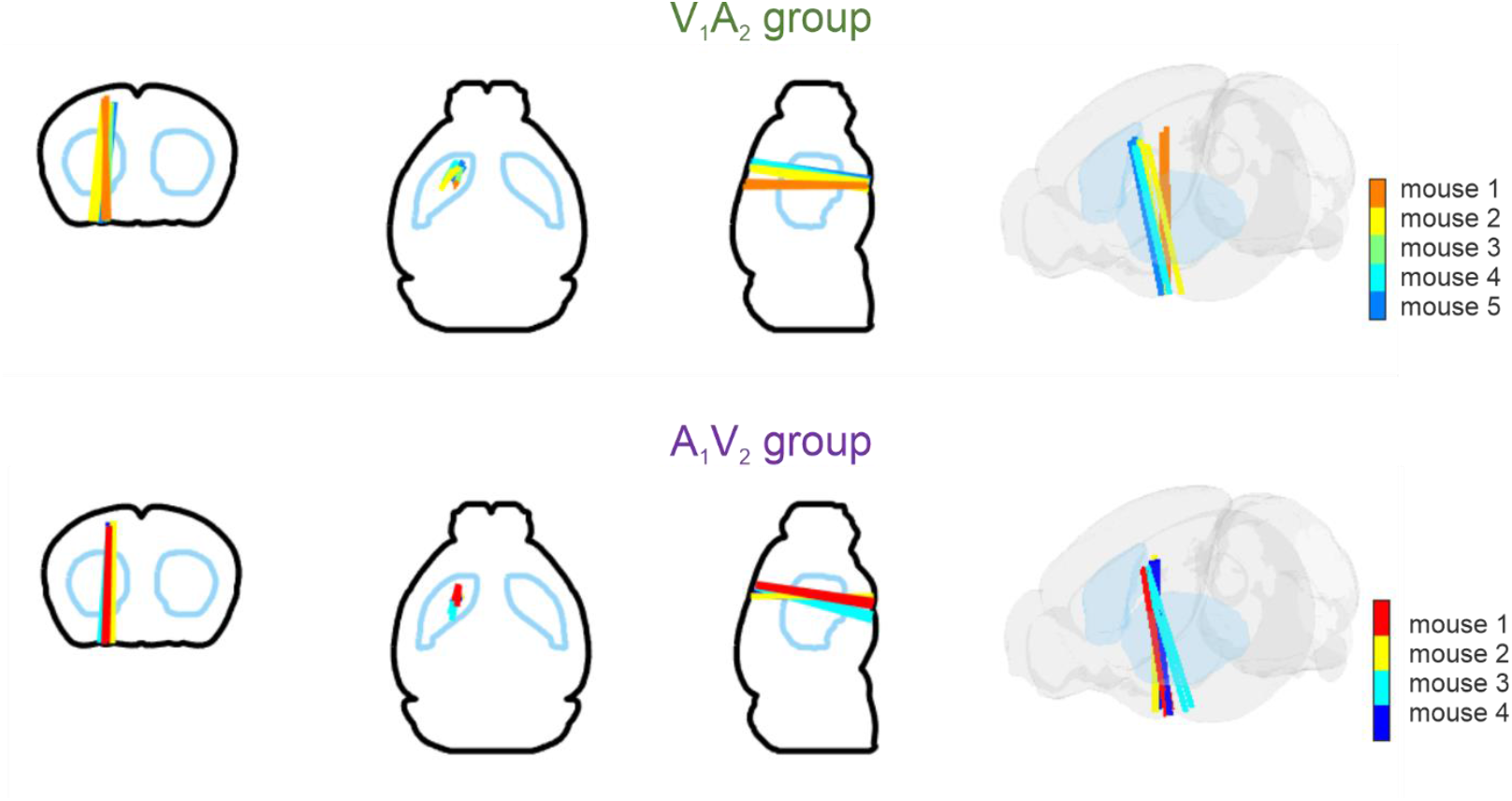
Probe positions in V_1_A_2_ and A_1_V_2_ mice. Schematic showing reconstructed Neuropixels probe tracks in V_1_A_2_ and A_1_V_2_ mice on the Allen CCF. Probe trajectories from all recordings are overlaid on coronal (left), axial (middle-left), sagittal (middle-right), and 3D views (right). Blue outline represents the striatum. Different colors represent individual mice, and different lines indicate different recording days. Recordings were in similar locations across both groups, indicating that observed differences in activity were not related to a difference in recording location.

